# CRISPR-mediated germline mutagenesis for genetic sterilization of *Anopheles gambiae* males

**DOI:** 10.1101/2023.06.13.544841

**Authors:** Andrea L. Smidler, Douglas G. Paton, George M. Church, Kevin M. Esvelt, W. Robert Shaw, Flaminia Catteruccia

**Affiliations:** Department of Immunology and Infectious Diseases, Harvard T.H. Chan School of Public Health, Boston, MA 02115, USA; Department of Genetics, Harvard Medical School, Boston, MA 02115, USA; Media Lab, Massachusetts Institute of Technology, Cambridge, MA 02139, USA; Howard Hughes Medical Institute, Chevy Chase MD 20815, USA

## Abstract

Rapid spread of insecticide resistance among anopheline mosquitoes threatens malaria elimination efforts, necessitating development of alternative vector control technologies. Sterile Insect Technique (SIT) has been successfully implemented in multiple insect pests to suppress field populations by the release of large numbers of sterile males, yet it has proven difficult to adapt to *Anopheles* vectors. Here we outline adaptation of a CRISPR-based genetic sterilization system to selectively ablate male sperm cells in the malaria mosquito *Anopheles gambiae*. We achieve robust mosaic biallelic mutagenesis of z*ero population growth* (*zpg*, a gene essential for differentiation of germ cells) in F1 individuals after intercrossing a germline-expressing Cas9 transgenic line to a line expressing *zpg*-targeting gRNAs. Approximately 95% of mutagenized males display complete genetic sterilization, and cause similarly high levels of infertility in their female mates. Using a fluorescence reporter that allows detection of the germline leads to a 100% accurate selection of spermless males, improving the system. These males cause a striking reduction in mosquito population size when released at field-like frequencies in competition cages against wild type males. These findings demonstrate that such a genetic system could be adopted for SIT against important malaria vectors.

## INTRODUCTION

Strategies aimed at targeting insect vectors of human pathogens are central to the control of vector-borne diseases and form a vital component of the WHO malaria control and elimination program (1). Increased implementation of vector control measures has contributed to a significant reduction in malaria-induced mortality rates (2), with the use of long lasting insecticide-treated nets (LLINs) and indoor residual spraying (IRS) contributing to over 75% of cases averted since the turn of the century (3, 4). However, these once-reliable control methods are becoming increasingly ineffective due to insecticide resistance mechanisms emerging in mosquito populations (5, 6), including resistance to all four classes of insecticides currently available for malaria control (7, 8), making the development of novel vector control technologies increasingly urgent.

Targeting insect reproduction has long proven an efficacious and sustainable approach for controlling and eradicating insect pests. One such technology, Sterile Insect Technique (SIT), relies on releasing large numbers of sterile male insects, inducing sterility in female mates and leading to a decline in the target insect population (9, 10). For SIT to be effective, sterile males need to be highly competitive against wild type males and effectively inhibit wild female remating (11). Traditionally, sterilization is achieved through irradiation or chemical-based sterilization methods to induce lethal DNA mutations in germ cells through oxidative stress (12). However, these methods of sterilization also impair overall male mating competitiveness: somatic DNA, lipid, and protein oxidation synergize to impact various life history traits (13), which combined severely reduce the male’s ability to compete for mates (14-18).

Developing sterilization methods that specifically target fertility genes may provide an alternative avenue to produce males that are fit for mating. Multiple, more precise, transgenic sterilization systems have been developed in some mosquito vectors, including those which preserve male fertility but kill offspring in post-embryonic developmental stages (19-22), those which express pro-apoptotic factors in the testes (23), and those which combine male sterilization and female-killing (24). While these systems cause transient species-specific population suppression following release, none have yet been adopted in the most important African malaria vector *Anopheles gambiae*. Fertility-reducing selfish genetic elements have been developed in this species using CRISPR/Cas technology (25, 26). These gene drive systems are very promising, although they can face rapid evolution of genetic resistance which hinders their application in the field (27). Importantly, the self-autonomous mode of propagation of gene drives necessitates safe mechanisms for containment and release which are not currently available (28). Malaria control would undoubtedly benefit from the development of alternative genetic sterilization systems that expand the genetic toolkit available to limit *An. gambiae* populations across Africa.

Similar to the precision-guided (pg) SIT system developed recently in *Drosophila melanogaster and Aedes aegypti* (24, 29), here we developed a safe, self-limiting and non-invasive CRISPR-based sterilization technology in *An. gambiae* that specifically disrupts a germ cell gene for SIT-based control of wild populations. Our target is *Zero Population Growth* (*zpg)*, a gap junction innexin which plays a crucial role in early germ cell differentiation and survival (30) and has been shown to be required for germ cell development in *Drosophila* (30, 31) and mosquitoes (32, 33). The *zpg* promoter has been demonstrated to express in a germline-specific manner (34), and in *An. gambiae zpg* knockdown by transient RNAi results in sterile males with phenotypically atrophied testes (32). Importantly, these males initiate typical post-mating responses in females following copulation and remain competent at mating, making *zpg* an ideal gene target for genetic sterilization. To generate sterile males, we developed a transgenic CRISPR system that achieves inducible mutation of *zpg* following a single cross of a germline-restricted Cas9-expressing line to a *zpg*-targeting gRNA-expressing line. We show that mosaic mutagenesis in the germlines of F1 males inheriting both transgenes is sufficient to achieve synchronous biallelic knockouts of *zpg* in the developing germline, ablating sperm development in 95% of males. Moreover, these males render females infertile after mating, and cause significant population suppression in competition cages against wild type males. With some adaptations, this system could be used for large-scale sterile male releases, providing a critical novel tool for self-limiting malaria vector control.

## RESULTS

### Male Δ*zpg* mosaics fail to develop normal testes

To generate spermless males, we crossed males expressing guide RNAs targeting *zpg* (gZPG line) to females expressing a germline-specific Cas9 (VZC line) (**Figure 1A**). (VZC/+; gZPG/+) offspring underwent significant mosaic mutagenesis in the germline, resulting in abnormal testes in the majority of males. This phenotype was robustly detectable from the pupal stage by the absence of fluorescence from a *Vas2*-EYFP reporter in the seventh abdominal segment (**Figure 1B**). Dissecting the reproductive tract from 126 adult males revealed atrophied testes with no visible mature sperm in 120 individuals (95.2%), in contrast to wild type controls (**Figure 1C, D**). A small minority of males showed however some level of germline differentiation and sperm development, having developed a single testis (5/126, 3.96%). A single male developed both testes (0.79%). In all 126 individuals, other reproductive tissues were unaffected, with male accessory glands appearing normal.

**Figure 1.**
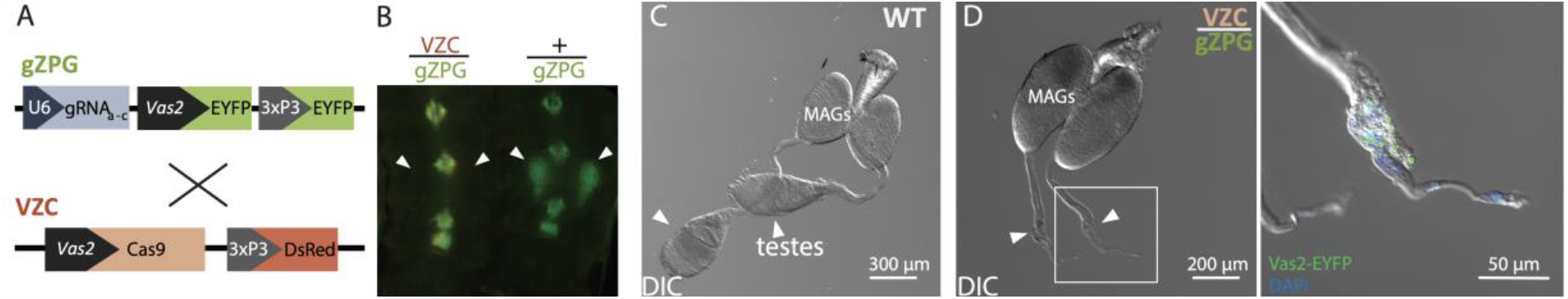
Crossing VZC and gZPG transgenic individuals generates spermless males. **A)** A schematic representation of the VZC and gZPG constructs used to generate (VZC/+; gZPG/+) males. These transgenic lines were previously described (35). In brief, VZC expresses Cas9 via the Vas2 promoter and carries a 3xP3-DsRed marker for selection. The transgene gZPG expresses three gRNAs; gRNA_a_, gRNA_b_ and gRNA_c_ in addition to a Vas2-EYFP germline marker and a 3xP3-EYFP selectable marker. Note the Vas2-EYFP fluorescent germline selectable marker that was used to screen for males with no clear evidence of sperm in their testes. **B)** Fluorescent testes can be observed through the pupal cuticle alongside the 3xP3-EYFP neural marker in gZPG males but not hybrid (VZC/+; gZPG/+) males. **C)** Wild type male reproductive tract showing male accessory glands (MAGs) and sperm-filled testes (arrowheads). **D)** In (VZC/+; gZPG/+) males, testes fail to develop (arrowheads), with minimal Vas2-EYFP and DAPI staining observed.

We sequenced the germline of some (VZC/+; gZPG/+) individuals and confirmed several CRISPR-induced mutations, mostly large deletions between the three gRNA target sites (**Figure 2A**), and some insertions (**Figure 2B**). Although this observation is qualitative, many of the large deletions observed appeared to result from mutagenesis under both gRNA_b,_ targeting the 3’ end, and gRNA_c_ at the 5’ end, with fewer initiated by gRNA_a_, suggesting differential cleavage capabilities of gRNA_c_ and gRNA_a_. Multiple mutations were observed within individual males (**Figure 2, sequences 7 & 10; 8 & 9**). Among the six males that showed some level of testis development, some sired progeny, and sequencing their testes revealed no evidence of mutagenesis (and their sequences are therefore omitted from **Figure 2B**). One male (**Figure 2B, Sequence 13)** instead harbored a 69 bp in-frame deletion roughly corresponding to the 4^th^ transmembrane domain of ZPG, suggesting sperm production can be maintained even in the presence of larger deletions. These data indicate that CRISPR mutagenesis of the male germline causes high levels of testis disruption but is not fully penetrant, and some fertility-maintaining mutations are possible.

**Figure 2.**
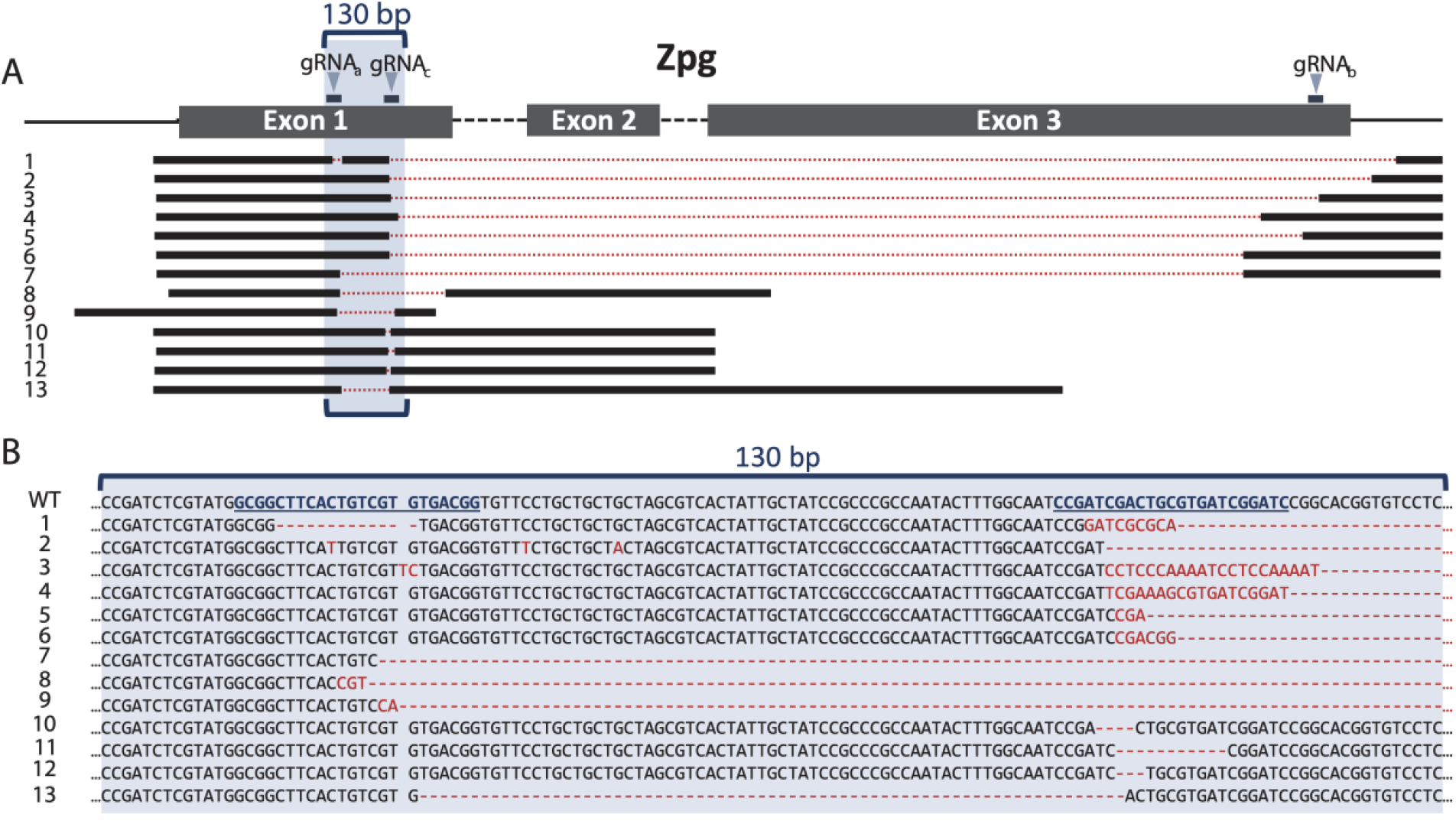
Germline CRISPR/Cas9 activity generates multiple large deletions in *zpg*. **A)** A representative map of observed mutations summarizing large deletions in the three exons of *zpg*. Positions of the three gRNAs used in this work are shown to scale. A 130 bp sequence encompassing gRNA_a_ and gRNA_c_ target sites is highlighted in blue. Sequence 13 belongs to a fertile male, while all others belong to sterile males. **B)** Sequences of observed mutations in the region between gRNA_a_ and gRNA_c_ (underlined). Sequences 1-13 correspond to 1-13 shown above in **A)**. Inserted bases are labelled in red and deleted regions are indicated by red dotted lines.

### Male Δ*zpg* mosaics are highly sterile

The absence of visible sperm in most (VZC/+; gZPG/+) males suggested that they should be sterile, making them good candidates for use in SIT programs. To test this, we released (VZC/+; gZPG/+) males into a cage with an excess of wild type ((+/+)) virgin females, and allowed them to mate for two nights. Females were then blood fed and allowed to lay eggs. Of the 4,132 eggs laid, only 3.05% were fertile, indicating high levels of sterility in females mated to (VZC/+; gZPG/+) males. To determine if hatched larvae were sired by a few fully fertile males or whether each male had some level of fertility, we performed individual forced mating assays between wild type females and (VZC/+; gZPG/+) males or wild type male controls, and assayed for fertility. While the vast majority of females mated to wild type males showed high fertility (more than 95%), females mated to (VZC/+; gZPG/+) males showed complete sterility in 25/26 cases (96%)(**Figure 3A**). The single female showing normal fertility levels produced a brood with an expected 50% (VZC/+): 50% (gZPG/+) transgene ratio. These results confirm that a minority of (VZC/+; gZPG/+) Δ*zpg* mosaic males maintain normal levels of fertility, likely due to failed mutagenesis or mutations that maintain fertility. Additional mating experiments using the parental (gZPG/gZPG) and (VZC/VZC) lines demonstrated that sterility is a product of *zpg* mutagenesis induced by the presence of both transgenes rather than non-specific effects of individual transgenes, as females mated to either (gZPG/gZPG) or (VZC/VZC) males had fertility levels comparable to females mated to wild types (**Figure 3B**).

**Figure 3.**
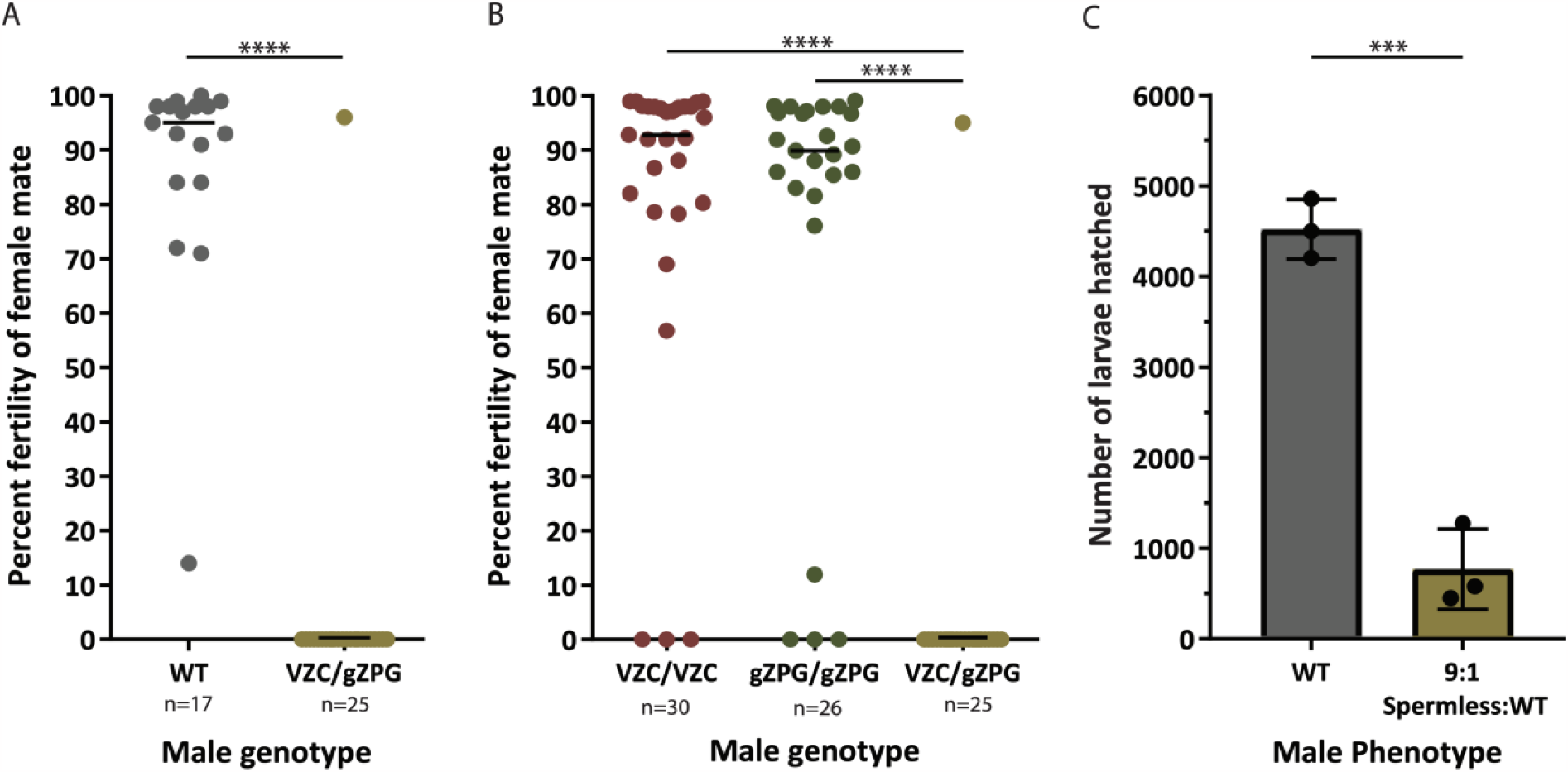
Δ*zpg* males are highly sterile and can suppress WT populations. **A)** Forced mating assays between WT female and either (VZC/+; gZPG/+) or WT males show most transgenic males are completely sterile (Mann Whitney, p < 0.0001). **B)** Forced mating assays between WT female and individual males of the (VZC/VZC), (gZPG/gZPG), or (VZC/+, gZPG/+) genotypes show the parental transgenes have no effects on male sterility (Kruskal-Wallis, p > 0.0001). **C)** Δ*zpg* males selected for lack of Vas2-EYFP fluorescence can effectively suppress numbers of larvae in competition cage experiments (Unpaired two-tailed t-test, p > 0.001).

### Male Δ*zpg* mosaics cause population suppression in cage releases

To be useful in SIT, genetically sterile males must be able to compete for female mates against field males. We tested whether (VZC/+; gZPG/+) males could suppress female fertility in competition with wild type males by simulating field releases in large cage assays. We used a 9:1 release ratio that is in line with ratios utilized in SIT strategies by introducing 90 (VZC/+; gZPG/+) males and 10 (+/+) males for three nights into cages containing 10 age-matched virgin females (9:1 Spermless:WT cages). For these experiments, we only selected males that showed no testes when analyzed by fluorescence, based on expression of the *Vas2*-EYFP germline marker. As control, we set up cages where only wild type males and females were introduced (WT cages). Following blood-feeding, in three replicate experiments we observed an 83% reduction in the number of larvae hatched in experimental cages compared to control cages **(Figure 3C**). Microscopic analysis of larvae from the experimental cages confirmed that none had been sired by transgenic males (0 out of 2305), suggesting these males are completely sterile. These results demonstrate that genetically sterile males maintain sufficient mating competitiveness to achieve significant population suppression in a competitive laboratory setting. Importantly, we observed only a negligible difference in wing length (a good proxy for male size, which is known to be linked to mating competitiveness (35)) between male groups that is unlikely to be biologically meaningful (Δ WT – (VZC/+; gZPG/+) = 46 ± 21 μm; p = 0.031, **Figure S1**).

## DISCUSSION

Generating sterile male *Anopheles* has historically faced developmental hurdles. Chemo-and radio-sterilization protocols have been developed (36), but generally cause a reduction in male competitiveness due to accumulated oxidative damage to cellular DNA, lipids and proteins (14-18, 37, 38). Moreover, chemical sterilization raises environmental concerns due to chemical residues after mass releases (39). GM technologies such as RIDL and pgSIT show great promise (19-21, 24) but have yet to be adopted in *An. gambiae*. Here we outline a system for generating genetically sterilized *An. gambiae* males that could be used in SIT-like programs against this important disease vector. We show that crosses between transgenic individuals expressing Cas9 in the germline and individuals expressing gRNAs targeting *zpg* efficiently produce sterile male F1 progeny. In the vast majority of cases, F1 males have atrophied testes, show no observable sperm, and harbor numerous CRISPR-generated mutant alleles that arise by active mosaic mutagenesis during development. When not pre-screened for testicular development by fluorescence, approximately 95% of these males completely sterilize their female mates, consistent with the penetrance of the mosaic spermless phenotype. We further demonstrate that removing males showing incomplete penetrance of the spermless phenotype by screening for *Vas2*-EYFP fluorescence at the pupal stage generates male populations that are completely sterile.

Anophelines are known to mate in large swarms with highly skewed sex ratios where competition between males is fierce (40). Competition cage assays with (VZC/+; gZPG/+) males show that transgenic spermless males can cause significant population suppression in the presence of wild type males, thereby demonstrating robust swarm fitness and mating competitiveness. Reduced mating competitiveness has often been observed with other sterilization methods. In the 1960’s and 1970’s chemo-sterilization was used to generate sterile males (41) but it exhibited peripheral mutagenic effects (39). Sterilization by radiation therefore became the dominant technique for most insects, and factors like age, stage, handling, oxygen level, ambient temperature and dose-rate were shown to be important to generate insects with sufficient competitiveness (42). In anophelines, irradiation at the adult stage, rather than the pupal stage, produces more competitive males (36, 37), but adult fitness is maximized only when a partially-sterilizing radiation dose is used, hindering suppression effects in trials (37). While males have similar longevity to wild type competitors (37), they nevertheless fail to compete for females, even when released in excess of modeled recommendations (38). Our results are compatible with previous studies that used RNA interference to knockdown *zpg* and that produced spermless males competent for mating (32), suggesting that specifically targeting this gene may not be harmful to males. It is possible the transgenes used here may impair male fitness in other ways, for example, through off-target mutagenesis of CRISPR/Cas9 activity (43). However, although obtained in limited laboratory conditions, our data show transgenic spermless males achieve significant population suppression in laboratory cages, indicating that their mutational loads do not significantly impair their mating competitiveness. In future studies, direct characterization of male mating competitiveness in semi-field settings will be critical to determine how this genetic sterilization system compares to traditional radiation-based sterilization techniques.

While our system shows promise for vector control, multiple steps of optimization will be required to render it functional in field settings. First, SIT strategies aim to release males that are >99% sterile, while we observed 5% of males escaping sterilization (10). To this end additional gRNAs could be used to boost genetic sterility but it will be important to understand the properties required for optimal DNA cleavage in the species. Others have shown that gRNAs vary in their mutagenic potential (44), an observation qualitatively supported by our findings where gRNA_C_ catalyzed more mutations than gRNA_a_. Alternatively, additional genes important for fertility could be targeted, such as those shown in *Drosophila* to be required in the germline, including *Tudor* (AGAP008268), *β2-tubulin* (AGAP008622), or *Vasa* (AGAP00857) among many possible candidates (reviewed in (45)). Optimization of the system to increase phenotype penetrance through genetic means, and/or addition of a fluorescent sorting step to remove partially sterile males would strongly improve the chance of successful suppression. Second, our system does not allow the automatic elimination of females from the released population, an essential requirement for any male release program (10). Combining genetic sterility with genetic sex separation systems such as those recently developed using CRISPR targeting of *femaleless* (46, 47) is therefore a necessary next step to operationalize genetic SIT for anopheline vectors.

A major hurdle facing successful development of genetic sterilization systems is that maintenance of highly sterile male-only lines is impossible by nature of their inability to breed. Therefore, all such systems require some degree of inducibility to suppress sterility until immediately prior to release. RIDL systems in *Aedes* achieve this by induction of a lethal transgene following release, which is suppressed by addition of tetracycline during rearing (20). In genetic SIT, because Δ*zpg* mosaic males and females are both infertile, inducibility is achieved by crossing two different transgenic populations, which alone do not show fertility defects. Although more cumbersome as two lines must be reared, this system facilitates mass rearing at scales sufficient for release. While this system requires significant optimization before it can be utilized in field settings, our work provides a valuable proof-of-principle that transgenic sterilization can enable SIT programs aimed at suppressing *Anopheles* populations.

Finally, it is important to note that, beyond its potential application for vector control, our system can be used to explore a variety of biological questions. Firstly, the role of sperm in regulating aspects of the female-post mating response is still largely unexplored. *An. gambiae* females display two major responses after copulation: the stimulation of oviposition following blood-feeding, and the induction of refractoriness to further mating. Both are initiated following sexual transfer of factors, including a male steroid hormone (48) from the male to the female atrium during copulation (48-50). Although a previous study showed that sperm is not involved in triggering these female responses (32), the use of transgenic spermless males may identify more subtle effects linked to sperm transfer and storage. Indeed, in *Drosophila*, sperm is needed to extend the mating refractoriness period up to a week by signaling through the slow release of male-transferred sex peptides bound to sperm tails (51-53). The Δ*zpg* mosaic males generated here could therefore be used to study the effect of sperm on similar post-mating responses in female mosquitoes, opening an intriguing avenue of research of significant importance for mosquito reproductive biology.

## METHODS

### Generating of transgenic mosquito lines

#### gRNA design

Design of gRNAs for these lines was previously reported in (54). Briefly, the *zpg* locus (AGAP 006241) was PCR amplified and sequenced across multiple individuals within our *An. gambiae* lines to identify any SNPs present. Putative gRNA candidates were identified by *in silico* tools available through the Broad Institute (https://portals.broadinstitute.org/gpp/public/analysis-tools/sgrna-design) and Zhang Laboratory at MIT (http://crispr.mit.edu (55)). Three gRNA targets were chosen to maximize the probability of mutagenesis early in the coding sequence, with the additional aim of achieving large deletions. Two gRNA candidates were chosen, gRNA_a_ and gRNA_c_, targeting the sequences (5’ GCGGCTTCACTGTCGTGTGACGG 3’) and (5’ CCGATCGACTGCGTGATCGGATC 3’) within Exon 1 located 71 bp and 150 bp from the stop codon respectively. They were further chosen for their localization over semi-unique restriction enzyme sites *Ale*I and *Pvu*I respectively to enable PCR-based identification of mutants, as previously described in (56). gRNA_b_ (5’ CCAAGTGTTTGCATTCCTGGCGG 3’) was designed to target the 3’UTR sequence to facilitate generation of large deletions. gRNAs under the control of the U6_57_ promoter (25) (composed of the 322 bp upstream of AGAP013557) were ordered as gBlocks (Integrated DNA Technologies, Skokie, IL) in two cassettes. gRNA_a_ and gRNA_b_ were synthesized as a tethered pair connected by a 21 bp sequence (5’ TTCACTGTGCGCATTATATAT 3’) predicted not to interfere with gRNA folding secondary structure (RNAfold, http://rna.tbi.univie.ac.at/cgi-bin/RNAWebSuite/RNAfold.cgi) (57). The gRNA_c_ was synthesized as an isolated gBlock.

### Plasmid Construction

As described previously (54), plasmids were constructed using standard molecular biological techniques and Golden Gate cloning (58, 59) into the *An. gambiae* transgenesis plasmids pDSAY and pDSAR (60).

#### VZC

To build VZC, the 2.3 kb *Vasa2* promoter (Vas2) (61) was PCR amplified from genomic DNA using primers (5’ CAGGTCTCAATCCCGATGTAGAACGCGAG 3’) and (5’ CGGTCTCACATATTGTTTCCTTTCTTTATTCACCGG 3’) and was cloned immediately upstream of SpCas9 amplified from plasmid PX165 (Addgene #48137) (62) using primers (5’ CAGGTCTCATATGGACTATAAGGACCACGACGGAG 3’) and (5’ CAGGTCTCAAAGCTTACTTTTTCTTTTTTGCCTGGCC 3’). These fragments were Golden Gate cloned into the multiple cloning site of the pDSAR vector, which provides an SV40 terminator for protein transcription termination, an attB site to facilitate ΦC31 transgenesis into well established *An. gambiae* transgenesis docking lines containing an attP, and a *3xP3*-DsRed fluorescence selectable marker (60). ***gZPG:*** To build gZPG, the two previously discussed gRNA-containing gBlocks were Golden Gate cloned into the multiple cloning site of the pDSAY transgenesis plasmid (60). To facilitate *in vivo* validation of the presence or absence of a germline, a *Vas2*-EYFP fluorescence cassette was further cloned into the unique *Asc*I site on the pDSAY plasmid backbone by Golden Gate ligation. For this cassette, the *Vas2* promoter was PCR amplified using the primers (5’ CGGTCTCACGCGCGATGTAGAACGCGAGCAAA 3’) and (5’ CGGTCTCACATATTGTTTCCTTTCTTTATTCACCGG 3’) and EYFP was PCR amplified with (5’ CAGGTCTCAATGGTGAGCAAGGGCG 3’) and (5’ CAGGTCTCAAAGCTTACTTGTACAGCTCGTCCATGCC 3’).

Complete plasmids were sequence verified by Psomagen Sequencing services (Rockville, MD, USA).

#### Transgenesis

Transgenesis procedures were carried out effectively as described in (54, 63, 64). The gZPG construct (350 ng/μl) was co-injected with a ΦC31-integrase expressing helper plasmid (80 ng/μl) into the posterior end of >3h-old aligned X13 docking line (60) *An. gambiae* embryos (n=1663), and the VasCas9 plasmid (350 ng/μl) was similarly injected into X1 docking line (60) embryos (n=2585). Survivors were reared to adulthood and outcrossed in bulk to large cages of wild-type *An. gambiae* G3 virgin adults (n>1000) of the opposite sex. New transformants were identified and isolated as newly hatched larvae in the subsequent F1 generation by fluorescence. F1 transformants were outcrossed to wild-type G3 to introduce genetic diversity before intercrossing in the subsequent F2 generation to establish homozygous lines. Homozygous lines were established by identification of homozygous individuals by fluorescence intensity and subsequent PCR verification.

### Generation of spermless (VZC/+; gZPG/+) males

To generate spermless males in bulk, (gZPG/gZPG) males were crossed to virgin (VZC/VZC) females in cages. Maternal deposition of Cas9 from VZC females facilitated increased mutagenic loads in the developing embryos leading to more penetrant mosaic phenotypes. Male pupae/adults were manually sex sorted from females under a microscope using a paintbrush. For forced mating experiments, spermless males were sex separated as pupae to guarantee virginity and their genotype was confirmed by dual (*3xP3*-EYFP; *3xP3*-DsRed) fluorescence. For caged competition experiments, male pupae were additionally screened for the absence of Vas2-EYFP from testicular tissues to remove males with an incompletely penetrant phenotype.

### Microscopy

Imaging of transgenic larvae and ventral pupal tails was carried out under a Leica M80 fluorescence dissecting microscope following immobilization on ice and positioning by paintbrush. Imaging of microscopic testes structure was carried out on a Zeiss Inverted Observer Z1 microscope following dissection in 1x PBS, and mounting in VECTASHIELD® Mounting Medium with DAPI within 1 h post-dissection. Tissues were dissected from 5-day-old virgin males.

### Mutation analysis

Male (VZC/+; gZPG/+) mutant testes or surviving larvae were analysed for mutations by PCR and sequenced for mutations. DNA extraction was carried out using the Qiagen DNeasy Blood & Tissue Kit, and PCR was carried out using a variety of primers flanking the *zpg* locus. Multiple primer pairs were used to capture large deletions and enable amplification over polymorphic regions. The forward primers (5’ CGTTTTCTTCACTCTCGGCACG 3’), (5’ GCAGCTTCTGGTAGTCGATGTCG 3’), and (5’ CCATTCGTTTGTTGCTGAAAGC 3’), and reverse primers (5’ GACCAGAAGCCGGAAAAGATC 3’), (5’ GAGGAACGCGGGTTTTTTTG 3’), and (5’ GTGAAATGTTTGGGCCCGATC 3’) were used in combinations to generate PCR products ranging from 700 bp to 5 kb. Individual mutant alleles were sequenced essentially as described in (56). PCR products were cloned into the CloneJet PCR Cloning Kit (ThermoFisher Scientific) to isolate PCR products corresponding to individual alleles, and plated on ampicillin (100 μg/mL) LB media plates. Individual colonies were either picked, cultured in liquid media, extracted (SpinSmart Plasmid Miniprep DNA Purification kit, Denville Scientific) and sequenced using the universal pJET2.1F or pJET2.1R primers (Psomagen USA), or the entire agar plate was sent for direct colony sequencing (Psomagen USA). Resulting sequencing reads were aligned to an annotated Snapgene 3.2.1 file of the *zpg* gene sequence.

### Infertility mating assays

#### Bulk mating

30 (VZC/+; gZPG/+) males were sexed as pupae and allowed to eclose into a 25 cm x 25 cm BugDorm cage (MegaView Science co, Taiwan). Four failed to eclose, leaving 26 surviving males for the experiment. Female pupae of the wild-type strain G3 were sexed on the same day and allowed to eclose in a separate cage. The absence of contaminating G3 males was confirmed the next morning, and 176 females were mouth-aspirated into the cage containing the (VZC/+; gZPG/+) males. Female mosquitoes were allowed to mate for 4 nights, and were blood fed on day 5 until significant diuresis was observed. An oviposition site consisting of a Whatman® filter paper cone (90mm, Grade 2, Sigma-Aldrich) within a urinalysis cup containing 80 ml deionised water was placed in the cage on day 7. The oviposition cup was removed on day 8, and larvae were counted and scored for transgene presence on day 9. Eggs and late-hatching larvae (none observed) were counted on day 11 and 12.

#### Individual forced-mating assays

5 days-post eclosion, virgin males of respective genotypes and blood-fed virgin wild-type G3 females were force-mated to guarantee paternity (method available at https://www.beiresources.org/MR4Home.aspx). Male carcasses were saved for subsequent mutation analysis. Successful mating was confirmed by autofluorescence of the mating plug in the female atrium, detectable through the female cuticle under a fluorescent microscope using a GFP filter set (previously demonstrated in (49)), and females were isolated to oviposit within individual paper cups lined with filter paper and filled with 1 cm deionised water. The number of eggs laid and larvae hatched were counted from each female’s brood, and larvae screened for transgene fluorescence to determine paternity. Escapee larvae sired by genetically sterilized (VZC/+; gZPG/+) fathers were collected for subsequent sequence analysis.

#### Cage competition assays

(VZC/+; gZPG/+) Vas2-EYFP-negative males and wild-type G3 males and females eclosed into separate cages, and adults were mixed at 3 days old to allow mating. Control cages contained 100 G3 males; 100 age-matched females, and competition cages contained 90 (VZC/+; gZPG/+) Vas2-EYFP-negative males and 10 G3 males; 100 G3 females. At 6 days old, females were offered a blood meal for 20 min and males were removed. An oviposition site was provided to females at 8 days old and was removed when 10 days old, and larvae were counted on days 10 and 11, and scored for genotype (and therefore paternity) by fluorescence.

## Respective Contributions

**ALS** conceived of the study and performed all cloning, transgenesis, mutant analysis, husbandry, data analysis, and drafted the manuscript. **ALS** and **KE** together performed transgene design. **ALS** and **WRS** executed the study and edited the manuscript. **DP** performed microscopy and edited the manuscript. **GMC** contributed to the study, provided CRISPR expertise and edited the manuscript. **FC** contributed to the design of the study, edited the manuscript, and provided entomological and reproductive expertise.

## Acknowledgements

A.S would like to sincerely thank Eryney Marrogi and Sean Scott for their research contributions and Emily Lund and Manuela Bernardi for their husbandry and administrative support respectively. This research was funded by the Howard Hughes Medical Institute/Bill and Melinda Gates Foundation grant OPP1158190 to F.C; by the National Institutes of Health (NIH) (award number R01 AI104956)to F.C.; and by an F31 AI120480-02 to A.S. F.C. is funded by the Howard Hughes Medical Institute (HHMI) as an HHMI investigator. Further funding was provided by Defense Advanced Research Projects Agency under the Safe Gene program to K.E. and G.C.; Burroughs Welcome Fund IRSA 1016432, and NIH R00-DK102669-04 to K.E. The findings and conclusions within this publication are those of the authors and do not necessarily reflect positions or policies of the HHMI or the NIH. The funders had no role in the study design, in data collection, analysis or interpretation, in the decision to publish, or the preparation of the manuscript.

