## Supplementary Data for "CRISPR-mediated germline mutagenesis for genetic sterilization of *Anopheles gambiae* males"

Figure S1

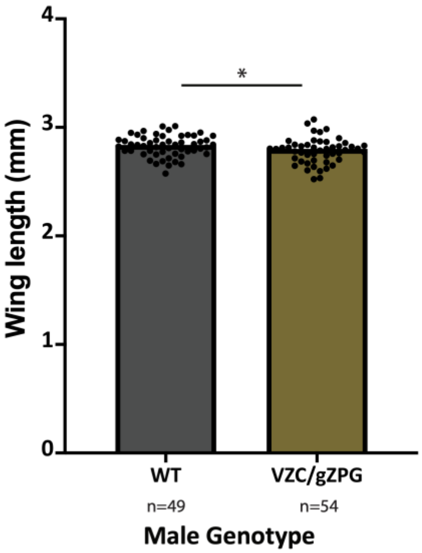


**Figure S1. Wing lengths of males used in mating competition assays.** Following the mating competition assays, males were removed, sorted by genotype by fluorescence, and their wings measured from the notch to the tip of the third cross vein using FIJI software. Average wing lengths differed slightly between the two groups, with wild type males begin marginally larger (Δ WT – {VZC/+; gZPG/+} = 46 ± 21 μm; $p = 0.031$).
